# Rapid and Live-cell Detection of Senescence in Mesenchymal Stem Cells by Micro Magnetic Resonance Relaxometry

**DOI:** 10.1101/2022.06.01.494362

**Authors:** Smitha Surendran Thamarath, Ching Ann Tee, Shu Hui Neo, Dahou Yang, Rashidah Othman, Laurie A. Boyer, Jongyoon Han

## Abstract

Detection of cellular senescence is important quality analytics for cell therapy products, including mesenchymal stromal cells (MSCs). However, their detection is critically limited by the lack of specific markers and the destructive assays used to read out these markers. Here, we establish a rapid, live-cell assay for detecting senescent cells using heterogeneous mesenchymal stromal cell (MSC) cultures. We report that the T_2_ relaxation time measured by microscale Magnetic Resonance Relaxometry (µMRR), which is related to intracellular iron accumulation, correlates strongly with senescent markers in MSC cultures under diverse conditions including different passages and donors, size-sorted MSCs by inertial spiral microfluidic device, and drug-induced senescence. In addition, the live-cell and non-destructive method presented here has general applicability to other cells and tissues, and can critically advance our understanding of cellular senescence.

## Introduction

Cellular senescence is a cell state that leads to cell cycle exit accompanied by genetic, metabolic, and morphological changes of cells due to aging and other external or internal conditions(Kuilman et al., 2010). Cellular senescence was described nearly 60 years ago based on the observation that cells in culture have a finite ability to divide.(HAYFLICK and MOORHEAD, 1961) Senescent cells are also found in tissues *in vivo*, and substantially implicated in many important pathologies. Cellular senescence is broadly defined as a complex cell fate that can be induced during normal development, physiological aging (longer time scales), or by intra- and extracellular stress (acute responses). Cellular senescence can be important for preventing cancer initiation and tumor progression(Campisi, 2013; Collado et al., 2007; Van Deursen, 2014; Hanahan and Weinberg, 2011). On the other hand, the accumulation of senescent cells in the body can have harmful effects, especially in age-related diseases like neurodegeneration, cardiovascular disease, osteoarthritis, renal dysfunction, non-alcoholic fatty liver disease, Type 2 diabetes, and cancer.(Campisi, 2013; McHugh and Gil, 2018; Naylor, R. M., Baker, D. J., van Deursen, 2013) The expression of genes involved in DNA replication, DNA repair, and cell cycle are often downregulated in senescent cells(Wagner et al., 2008). In addition, senescent cells display a senescence-associated secretory phenotype (SASP), which causes an alteration in the tissue microenvironment, including local or systemic inflammation and disruption of normal tissue structure, leading to resistance to immune clearance of these pathological cells(Campisi, 2013; Mareschi et al., 2006; Naylor, R. M., Baker, D. J., van Deursen, 2013). Thus, the detection and elimination of cells with senescent characteristics have become a target for medical intervention (Van Deursen, 2014).

Detecting and quantifying senescent cells is especially critical in cell therapy bioproduction (Bonab et al., 2006). For example, mesenchymal stem/stromal cells (MSCs) have clinical applications for indications in ischemic, inflammatory, autoimmune, regenerative potential, and degenerative disorders(Mareschi et al., 2006). Yet, senescent MSCs impose significant challenges in their clinical applications (Bonab et al., 2006; Drela et al., 2019). Despite the clinical potential of MSCs, proliferation arrest is observed in MSC populations, along with morphological and phenotype alterations, after prolonged cultureor expansion *in vitro* (Banfi et al., 2000; Bonab et al., 2006; Rubin, 2002; Schellenberg et al., 2011). Senescent MSCs exhibit enlarged and flat cell morphologies with stiffened nuclei and granular cytoplasm(Bonab et al., 2006; Campisi, 2013; Li et al., 2017; Schellenberg et al., 2011). Senescent MSCs have decreased multilineage differentiation potential(Banfi et al., 2000; Bonab et al., 2006; Noer et al., 2007; Schellenberg et al., 2011) and altered secretory and immunomodulatory functions, resulting in reduced therapeutic value(Campioni et al., 2009; Campisi, 2013; Liang et al., 2013). Notably, MSCs isolated from older donors show a higher level of senescence than younger donors (Zhou et al., 2008). Therefore, it is crucial to determine the senescent state of MSCs and other cell-based products as part of quality control in therapeutic cell production.

A critical bottleneck is the lack of quantitative tools that agnostically detect senescent MSCs in heterogeneous cell populations. Phenotypic diversity of senescent cells and a lack of robust biomarkers have hampered progress in both our understanding of cellular senescence and therapeutic applications.(Zhang et al., 2021),(Mehdizadeh et al., 2021) The standard method for detecting senescent MSCs is histochemical staining of acidic lysosomal β-galactosidase (β-gal)(Campisi, 2013; Dimri et al., 1995). The staining procedure requires stable pH (∼6), cell fixation, and a long incubation period (>12 hours). Alternatively, a modified colony-forming unit assay (CFU-f) can detect cell senescence by estimating the proliferative potential of the MSCs after multiple passages(Penfornis and Pochampally, 2016; Schellenberg, A., Hemeda, H. & Wagner, 2013; Schellenberg et al., 2011). Real-time polymerase chain reaction (RT-qPCR) can also detect senescence by measuring expression levels of senescence-associated markers p16, p21, p53(Campisi, 2013; Cheng et al., 2011; Naylor, R. M., Baker, D. J., van Deursen, 2013). However, these assays require lengthy and laborious procedures and are destructive end-point assays, that are not adequate for quality control of MSCs during cell manufacturing (Debacq-Chainiaux et al., 2009). In addition,. Therefore, a rapid detection method that allows real-time quantification of senescent MSCs is critical for the quality control of cell therapeutics for regenerative medicine.

Significant evidence indicates that iron homeostasis and its dysfunction correlate with aging and related pathologies(Fairweather-Tait et al., 2014; Morris et al., 1986; Ogilvie-Harris and Fornaiser, 1980; Puxeddu et al., 2014; Xu et al., 2012). Cellular iron is present in Fe^2+^ (reactive, labile, or ‘free’ iron, diamagnetic) or Fe^3+^ (paramagnetic, often bound with iron storage protein ferritin) forms (**Figure 1a**). In iron-homeostatic cells, uptake, balancing, and recycling of Fe^2+^/Fe^3+^ are maintained by various iron transport system proteins (*e*.*g*., transferrin (Tf), TfR1, ferroportin, NCOA4, DMT1) and regulators (*e*.*g*., p53, NRF2)(Chen et al., 2020), involving many different molecular pathways(Zhou et al., 2020). Notably, growing evidence indicates that an increase in intracellular iron may serve as a ‘natural marker’ for cellular senescence. Senescent cells accumulate iron (ferritin-bound Fe^3+^) up to ∼30 fold by adjusting their iron homeostasis proteins(Masaldan et al., 2018) and inhibiting ferrinitophagy(Mancias, J. D., Wang, X., Gygi, S. P., Harper, J. W., Kimmelman, 2014) (autophagy of Fe^3+^-bound ferritin) that recycles stored iron back to Fe^2+^. This sequestration of iron in ferritin (Fe^3+^) prevents ferroptosis(Dixon et al., 2012), a unique cell death pathway requiring labile iron (Fe^2+^) released from ferritinophagy(Gao et al., 2016). Therefore, directly quantifying intracellular iron (Fe^3+^) in live cells could provide a measure for detecting cellular senescencein real time during bioproduction.

**Figure 1.**
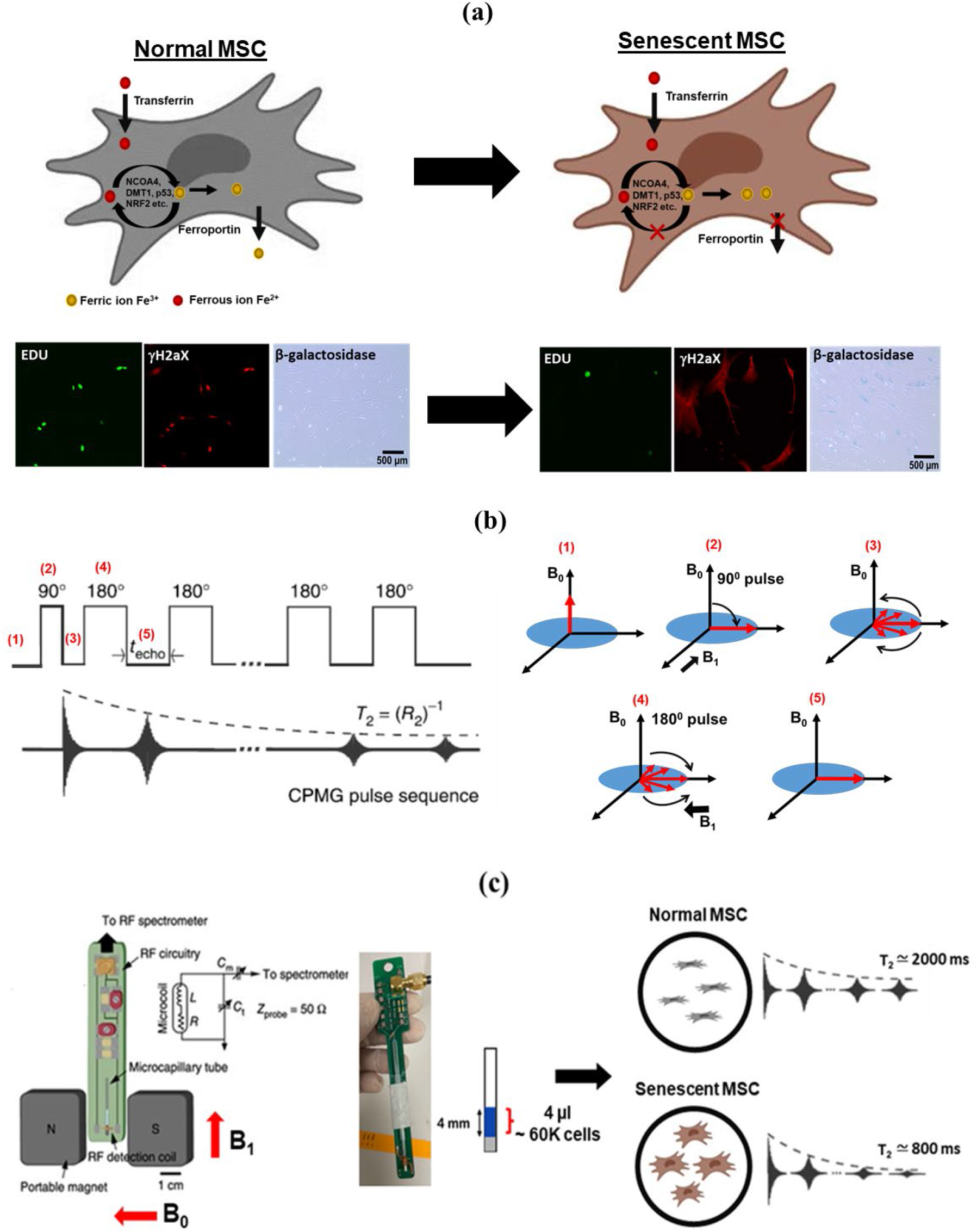
Live-cell and rapid detection of senescent MSCs using Magnetic Resonance Relaxometry. (a) Normal cells are maintaining iron homeostasis, mediated by numerous iron transporters and iron-binding proteins. Senescent cells are correlated with the accumulation of paramagnetic Fe^3+^ (an inactive form of iron). Normal cells and senescent cells are analyzed by standard assays are shown in the below images. The cells are stained by the senescence marker (γ-H2aX foci, red fluorescence), the proliferation marker, 5-ethynyl-2′-deoxyuridine (Edu, green fluorescence), and β-galactosidase staining (stain the cytoplasm of senescent cells as blue). (b) The CPMG (Carr-Purcell-Meiboom-Gill) pulse sequence for measuring the T_2_ values, which works efficiently in inhomogeneous magnetic fields produced by the permanent magnet of µMRR. A train of radiofrequency pulses is applied to the proton nuclei at the resonance frequency of 21.65 MHz with the inter echo time interval *t*_echo_. It is repeated for thousands of echoes until an equilibrium is reached. This decaying peak height of successive echoes over time is called transverse relaxation time T_2_. (c) MRR system consists of a permanent magnet that provides a strong magnetic field (B_0_). A home-built radiofrequency (RF) detection probe is connected to an RF spectrometer. The MSC sample (young/proliferating/senescent MSCs) in the microcapillary tube is placed in the RF detection coil for T_2_ measurements. The microcapillary tube contains the 4 µl of MSC sample (blue) in the 4 mm detection range of the RF detection coil. The typical cell number required is 60,000 MSCs within the detection volume. The grey color is the crystoseal to seal the microcapillary tube. The right side shows the ^1^H spin-spin relaxation time T_2_ of normal and senescent MSCs in which decreased T_2_ in senescent MSCs than normal MSCs.

We previously reported microscale-Magnetic Resonance Relaxometry (µMRR) as an efficient malaria diagnostic(Peng et al., 2014). The increased magnetic susceptibility of paramagnetic Fe^3+^ in the hemozoin crystals of infected red blood cells (RBCs) caused faster transverse relaxation of protons (T_2_) than the diamagnetic Fe^2+^ state in uninfected/healthy RBCs(Peng et al., 2014),(Thamarath et al., 2019). In addition, µMRR-based phenotyping of the oxidative stress response in diabetes mellitus patients’ blood was reported as a potential alternative to the conventional Hb-A1c test(Peng et al., 2020). Others also reported the high-sensitivity detection of tumor cells(Castro; Haun, J. B., Devaraj, N. K., Hilderbrand, S. A., Lee, H. & Weissleder, 2010; Lee, H., Sun, E., Ham, D. & Weissleder, 2008), bacteria(Liong et al., 2013), and tuberculosis(Issadore, 2011) using a similar µMRR device with immunomagnetic labeling of molecular and cellular targets. Here, we show that µMRR is a non-destructive, label-free, and rapid method for detecting senescence in heterogeneous MSC cultures over a range of conditions. Thus, µMRR represents a robust methodology for quality control for cell therapy manufacturing, opening the door for broader adaptation of this method for detecting cellular senescence in other cells and tissues.

## Results

### MRR detection of senescent MSCs and its correlation with standard assays

Senescent cells in MSC cultures often diminish their therapeutic efficacy in preclinical and clinical trials. We sought to measure proliferation and senescence using µMRR by comparison with conventional assays to correlate cell state directly with the T_2_ values. The CPMG (Carr-Purcell-Meiboom-Gill) pulse sequence (**Figure 1b**) is used for measuring the T_2_ values, which works efficiently in inhomogeneous magnetic fields produced by a permanent magnet of µMRR. The µMRR system (**Figure 1C**) consists of a 0.5 Tesla permanent portable magnet and a radio frequency (RF) detection probe connected to a spectrometer. For MRR experiments, a small-sized sample (4µl sample volume, which fits into the detection range of RF detection coil) is required, which is loaded into the microcapillary tube with minimal processing. T_2_ values of the samples having 6×10^4^ cells are measured within 5 minutes without damaging the cells (**Figure S1**).

For comparison between conventional senescence assays and µMRR, we cultured MSCs *in vitro* from the same donor to passage 3 (P3), passage 5 (P5), and passage 6 (P6). As a positive control for senescence cells, we also treated cells from P3 for 24 hours with 1µM of doxorubicin (DOX), a chemotherapy agent that produces DNA damage and reactive oxygen species known to induce senescence in tumor cells.(Kozhukharova et al., 2018) To measure DNA synthesis, EdU (5-ethynyl-2 ’ - deoxyuridine) was used as an indicator of active cell proliferation (shown in green). In contrast, senescent cells are known to accumulate γ-H2AX foci (indicated in red). A proliferation (Pf) index and senescence (Sn) index were calculated for all passages and DOX-treated MSCs from these images (see Methods and materials) (**Figure 2a**) and plotted respectively (**Figure 2b and 2c**). Based on the Pf/Sn index graphs, more proliferating cells (EDU) were present in P3. Pf gradually decreased with the later passages (P5 and P6) and was lowest in DOX-treated cells. The Sn Index showed the opposite trend, where more senescent-like cells (γ-H2aX foci) were observed in P6 and DOX-treated cells than P3 and P5. Moreover, we observed a higher proportion of β-gal-stained cells in later passages (P6) (**Figure 2d**). Similarly, qPCR of the senescence markers p16 and p21 showed higher expression in P6 (**Figure 2e**). Thus, senescent MSCs were clearly visible at later passages and in response to DOX treatment.

**Figure 2.**
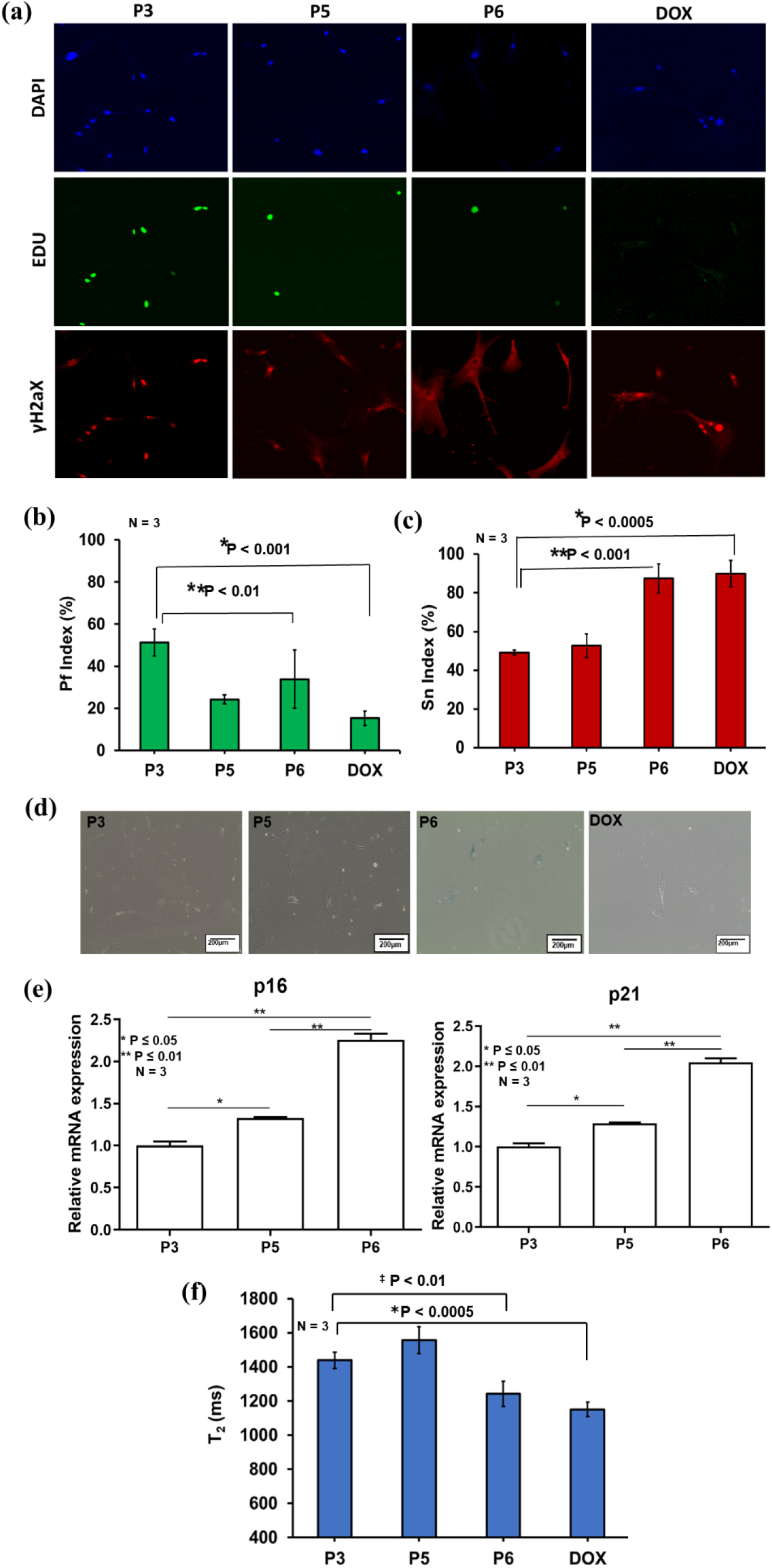
Correlating MRR with standard assays. (a) Immunofluorescent images of MSCs that are culture-expanded to passages P3, P5, P6, and doxorubicin treated MSCs from P3. The cells are stained by the senescence marker (γ-H2aX foci, red fluorescence) and the proliferation marker, 5-ethynyl-2′-deoxyuridine (Edu, green fluorescence). The cell nuclei are counterstained with NucBlue (Hoechst 33342), which emits blue fluorescence (shown as DAPI) when bound to DNA. The experiment was performed with 3 technical replicates and more than 100 cells in each replicates. (b & c) The proliferation (Pf) Index (%) and Senescent (Sn) Index (%) are calculated by counting number the green and red fluorescent stains relative to the number of blue fluorescent stains in all images. A total of 10 images from each of 3 technical replicates of staining experiments in MSCs of P3, P5, P6, and DOX treated were used for the calculation of Pf Index (%) and Sn Index (%), which are plotted in (b) and (c) respectively. Statistical analyses of P3 with DOX and P6 (n=3) are done by a two-tailed T-Test. For (b) the P values are *P < 0.001 and ^‡^P<0.01. For (c) the P values are *P < 0.0005 and ^‡^P < 0.001. (d) β-galactosidase staining of MSCs for passages P3, P5, and P6 where the cytoplasm of senescent MSCs was stained as blue. (e) Relative mRNA expression values of senescence-associated markers p16 and p21 for MSCs from a single donor at different passages (P) namely P3, P5, and P6. Data shown is representative of 3 technical replicates for each passage. Unpaired two-tailed t-test was performed between selected pairs to determine the statistical significance. * represents P≤0.05, ** represents P≤0.01. (f) Average T_2_ results (n=3) of P3, P5, P6 and DOX treated. Statistical analysis of P3 with DOX and P6 (n=3) are done by 2-tailed T-test and the P values are *P < 0.0005 and ^‡^P < 0.01.

We next compared these data with T_2_ measurements by performing parallel µMRR experiments using a normalized cell number of 6 ×10^4^ in 4µl volume in the microcapillary tube. MSCs were added to the micro-capillary tube and placed inside the radio frequency (RF) detection coil of the µMRR machine. We demonstrate that the T_2_ measurements correlated well with results from conventional end-point assays. Specifically, we observed a significant decrease in T_2_ levels, an indicator of increased iron accumulation, at later passages (P6) and after DOX treatment (**Figure 2f**). Thus, MRR T_2_ measurements correlate well with known biochemical senescence markers.

### MRR detection of induced senescence by treatment of cytokines

Cytokines such as Transforming Growth Factor-β1 (TGFβ1) and interleukin −1 (IL-1) induce senescence in MSCs.(Tominaga and Suzuki, 2019) TGFβ1 is a TGF family member that controls numerous cellular functions, including proliferation and cell death(Ksiązek, 2009; Wu, J., Niu, J., Li, X., Wang, X., Guo, Z., Zhang, 2014). IL-1 is a pro-inflammatory cytokine whose expression is associated with SASP(Lunyak et al., 2017). We next treated MSCs (same donor, passage 6) with different concentrations (10ng and 20ng/ml) of TGFβ1 and IL-1 for 3 days and performed measurements using µMRR. T_2_ values of TGFβ1 and IL-1 treated cells decreased with increasing cytokine concentrations (**Figure 3a**), predicting that TGFβ1 and IL-1 induce MSC senescence in culture. Quantification of β-galactosidase staining shows higher levels of senescent cells in TGF-β1 compared to IL-1 treated cells (see **Figure 3b**), in accordance with T_2_ values.

**Figure 3.**
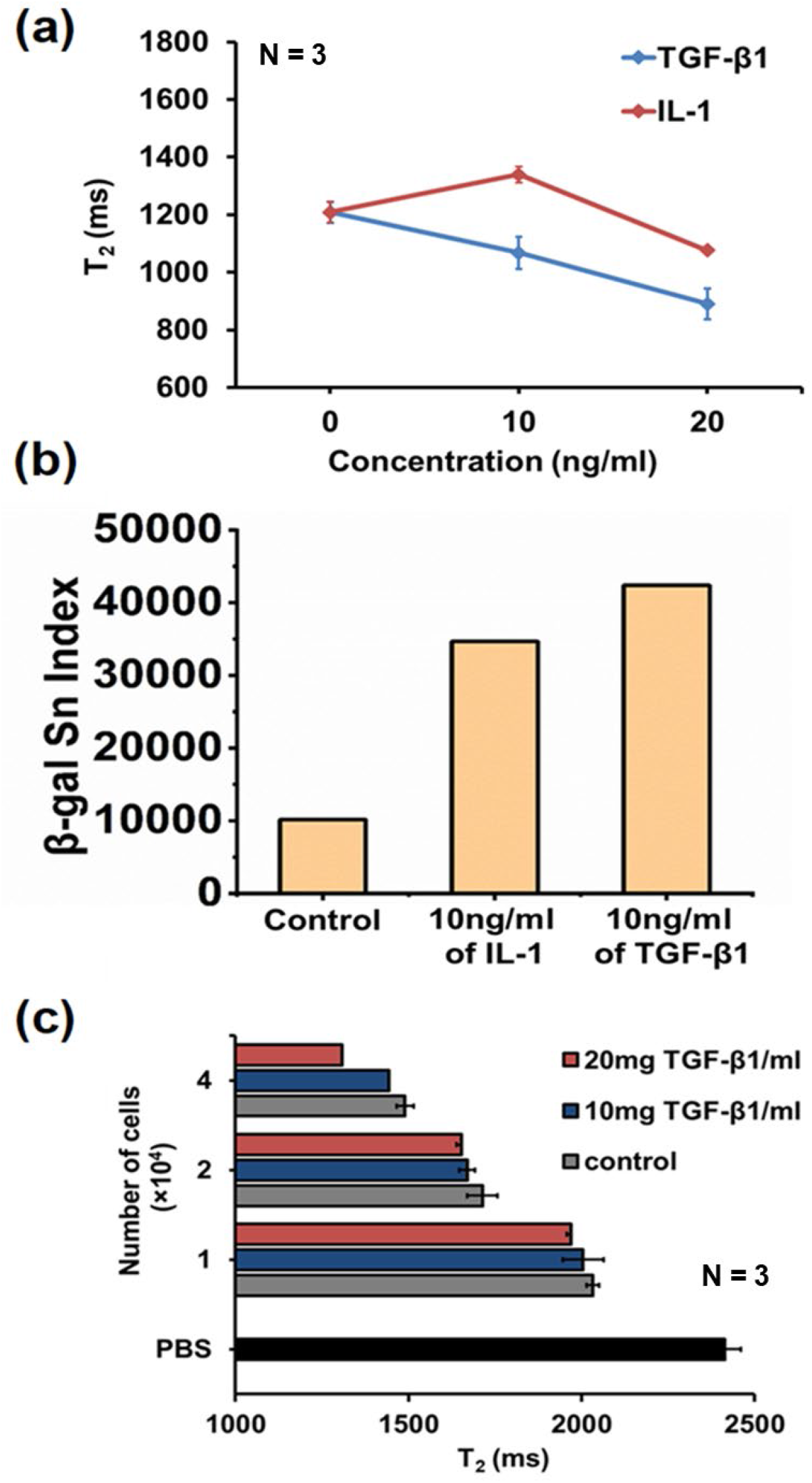
MRR detection of induced senescence by treatment of cytokines. (a) Average T_2_ values (n=3) of MSCs (same donor, Passage 4) control and treated with TGB-β1 (blue) and IL-1 (red) at different concentrations (10 and 20 mg/ml). (b) Quantification of Senescence Index from β-galactosidase staining images of MSCs control (a), 10 ng/ml of IL-1 treated MSCs (b) and 10 ng/ml of TGF-β1 treated MSCs (c). The image of each sample contains more than 200 cells. (c) Limit of detection of MRR assay for senescent MSCs. Average T_2_ values (n=3) of MSCs with increasing cell numbers in microcapillary tube of MRR for control and TGF-β1 (10 and 20 mg/ml) treated MSCs of the same donor at passage 4. The T_2_ value (n=3) of PBS in the microcapillary tube is shown as a black bar at the bottom of figure (b).

Since MSCs donors typically show variability in proliferative ability, it is important to find the limit of detection (LOD) of the µMRR assay in terms of the cell number. Thus, we analyzed the LOD by titirating the number of cells to 4 ×10^4^, 2×10^4^, and 1×10^4^ cells for µMRR detection as shown in **Figure 3c and Figure S2**. MSCs (same donor, P6) were treated with TGFβ1 at different concentrations (10 and 20mg/ml) to induce senescence, and batches of 1, 2, and 4 (×10^4^ cells) in 4µl detection volume of TGF-β1 treated and untreated MSCs were collected after 3 days for µMRR detection. The T_2_ value of PBS buffer was also compared with T_2_ value of MSCs (treated and untreated) as a control. This analysis shows a significant difference in T_2_ value of TGFβ1 treated MSCs when compared to the T_2_ value of untreated MSCs and PBS at cell numbers at or larger than 4×10^4^ cells. This establishes the cell number necessary for a reliable µMRR quantification of senescent cells in live cultures, and we used 6×10^4^ cells per assay in all of our experiments.

### MRR detects increase in senescence of large MSCs enriched by spiral microfluidic device

Previously, we and others identified cell size as a marker for distinguishing MSC phenotypes(Colter et al., 2001; Lee et al., 2014). Smaller MSCs are generally multipotent MSCs, whereas larger MSCs tend to display limited growth and differentiate more prominently toward the bone lineage. We found that large MSCs sorted by inertial microfluidic devices displayed senescent phenotypes with limited growth potential and longer doubling times compared to smaller MSCs(Yin et al., 2018) (**Figure 4a**). Using the same microfluidic device, we sorted MSCs into three size groups: Unsorted, Small (11-15 µm), Medium (15-22 µm), and Large (22-26 µm) (**Figure 4b and Figure S3**). Analysis of the sorted MSCs showed decreasing T_2_ value from Small to Large cells (**Figure 4c**), indicating water proton nuclear relaxation, which is ∼2000ms in pure water, is significantly reduced due to the paramagnetic iron (Fe^3+^) impurity in senescent MSCs. In addition, expression of senescence-associated markers p16 and p21 (measured by RT-qPCR) were elevated in the Large cells compared to Medium and Small groups (**Figure 4d**). We also observed a high β-gal Sn Index in Large and Unsorted group cells (Figure 4e and Figure S4), in line with the lower T_2_ values measured by µMRR. These results support previous reports of Fe^3+^ storage in senescent cells(Masaldan et al., 2018), because faster T_2_ relaxation is due to increased paramagnetic Fe^3+^ content in the cells. We also measured the secretion of molecules associated with the senescence-associated secretory phenotype (SASP) for the unsorted and size-sorted MSC subpopulations (Small, Medium, and Large) (**Figure 3f**). Using Luminex assays, we found higher expression of key secreted markers associated with SASP such as MCP-1, IL-6, IL-8(Suvakov et al., 2019),(Soto-Gamez and Demaria, 2017), and TGF-β1(Tominaga and Suzuki, 2019) in larger MSCs(Yin et al., 2020), consistent with the decrease in T_2_ values.

### MRR-based measurements correlate with the multilineage differentiation potential of MSCs

The most critical therapeutic quality attribute of MSCs is their ability to differentiate into cell lineages such as adipocytes, chondrocytes, and osteoblasts. MSCs isolated from different donors vary significantly in their multilineage differentiation potential, a critical and poorly understood limitation in achieving consistent quality control of MSCs for therapy. Here, we analyzed MSCs from five different donors (D1-D5; See Methods for description of donors and Table 2) cultured *in vitro* to P3. For µMRR analysis, we used a normalized cell number of 6×10^4^ for each T_2_ experiment. **Figure 5a** shows the T_2_ profile of MSCs from different donors. Multilineage differentiation potential of MSCs from the corresponding donors (D1-D5) was then measured by staining Alizarin red S, Oil Red O, and Safranin O for osteogenesis, adipogenesis, and chondrogenesis, respectively (**Figure S5**). The multilineage differentiation index was calculated (**Figure 5b and 5c**), and T_2_ values of MSCs from each donor were correlated with differentiation potential. Chondrogenesis Index correlated with the T_2_ value, where the highest T_2_ value was recorded for MSCs from D1 and the lowest T_2_ from D5. The population doubling time in days for each passage of donors D3 and D4 were also measured (**Figure S5**), which correlates with the T_2_ profile of respective donors. Hence, the lower T_2_ values corresponded to a decline in chondrogenic differentiation potential and reduced proliferation. Thus, decreased differentiation potential in MSCs from certain donors may be affected by the presence of higher proportions of senescent cells in the population, and this could also influence cell proliferation.

**Figure 4.**
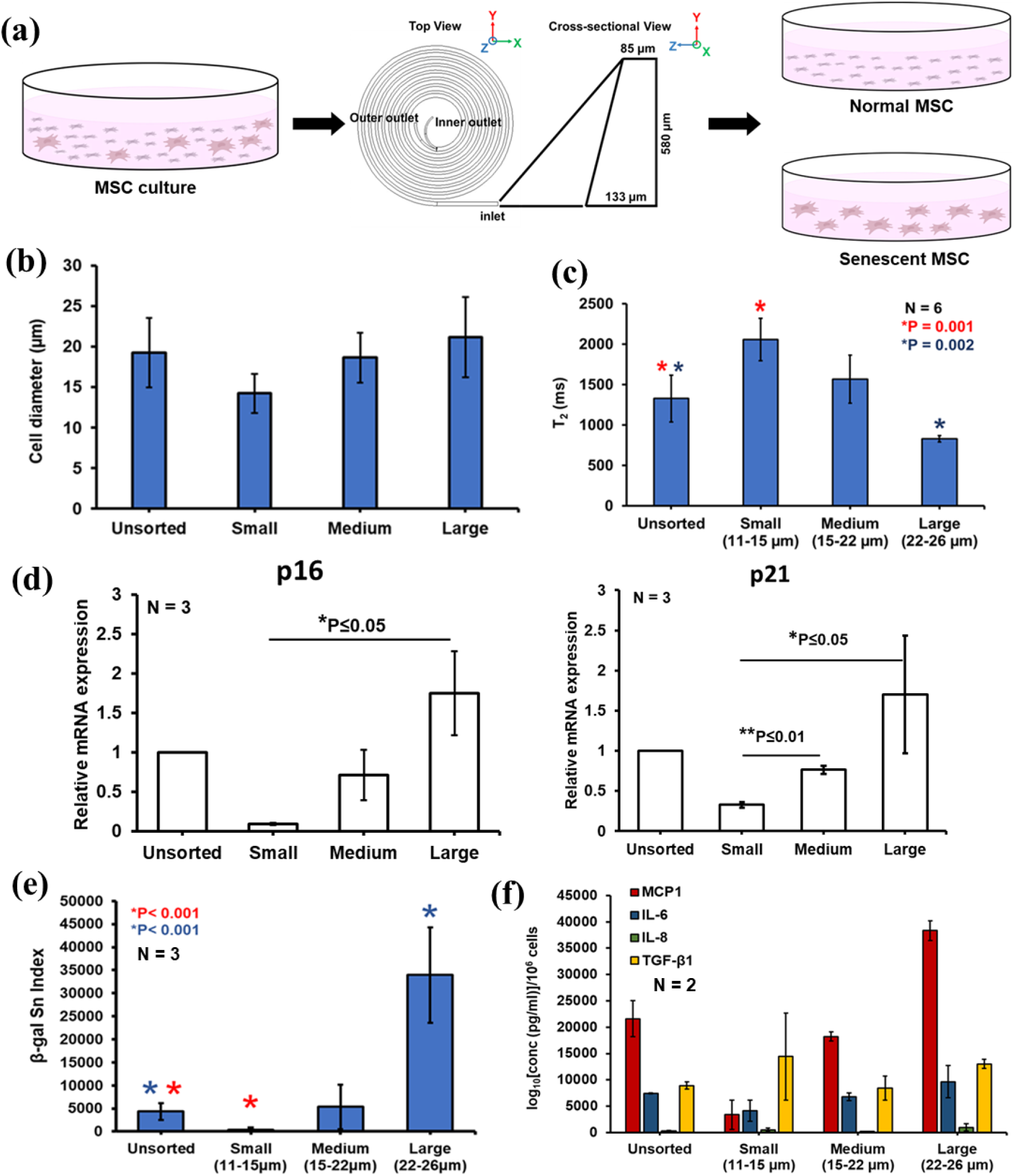
MRR assay shows large MSCs are more senescent based on size sorting using a spiral microfluidic device. (a) Separation of young/proliferating and senescent MSCs using a spiral microfluidic device from MSC culture expansion. The MSC culture expansion is pumped into the microfluidic device and sorted at various speeds to collect different sized MSCs at the outlets. (b) Cell diameter (µm) of sorted MSCs using spiral microfluidic sorting device. (c) Average T_2_ values (n=6) of unsorted and size sorted MSCs from two different donors using a spiral microfluidic device. Large cells (22-26 µm) are the size sorted MSCs collected in the first round of sorting, and Medium (15-22 µm) & Small (11-15 µm) cells are collected from the 2^nd^ round of sorting. Statistical analysis of unsorted & small cells (n=6, *P=0.001) and unsorted & Large cells (n=6, *P=0.002) are done by unpaired two-tailed t-test. (d) mRNA expression level of unsorted and size-sorted MSCs showing the overexpression of senescent markers P16 and P21 in LargeMSCs. Data shown is representative of 3 technical replicates for each group. Unpaired two-tailed t-test was performed between selected pairs to determine the statistical significance. * represents P≤0.05, ** represents P≤0.01 (e) Quantification of Senescence Index from the β-galactosidase images of unsorted and size sorted MSCs. Five to ten random areas were captured at the bright field by a color camera for quantifying the β-galactosidase staining-based senescence index (β-gal Sn Index) using MATLAB image processing. Biological replicates N=3. The images of each sample contains more than 500 cells. (f) Secretome profiling of factors associated with senescence secreted from unsorted and size-sorted MSC subpopulations in two different donors. Each sample was performed in technical duplicates. Concentrations of the secreted analytes normalized per million cells were plotted.

**Figure 5.**
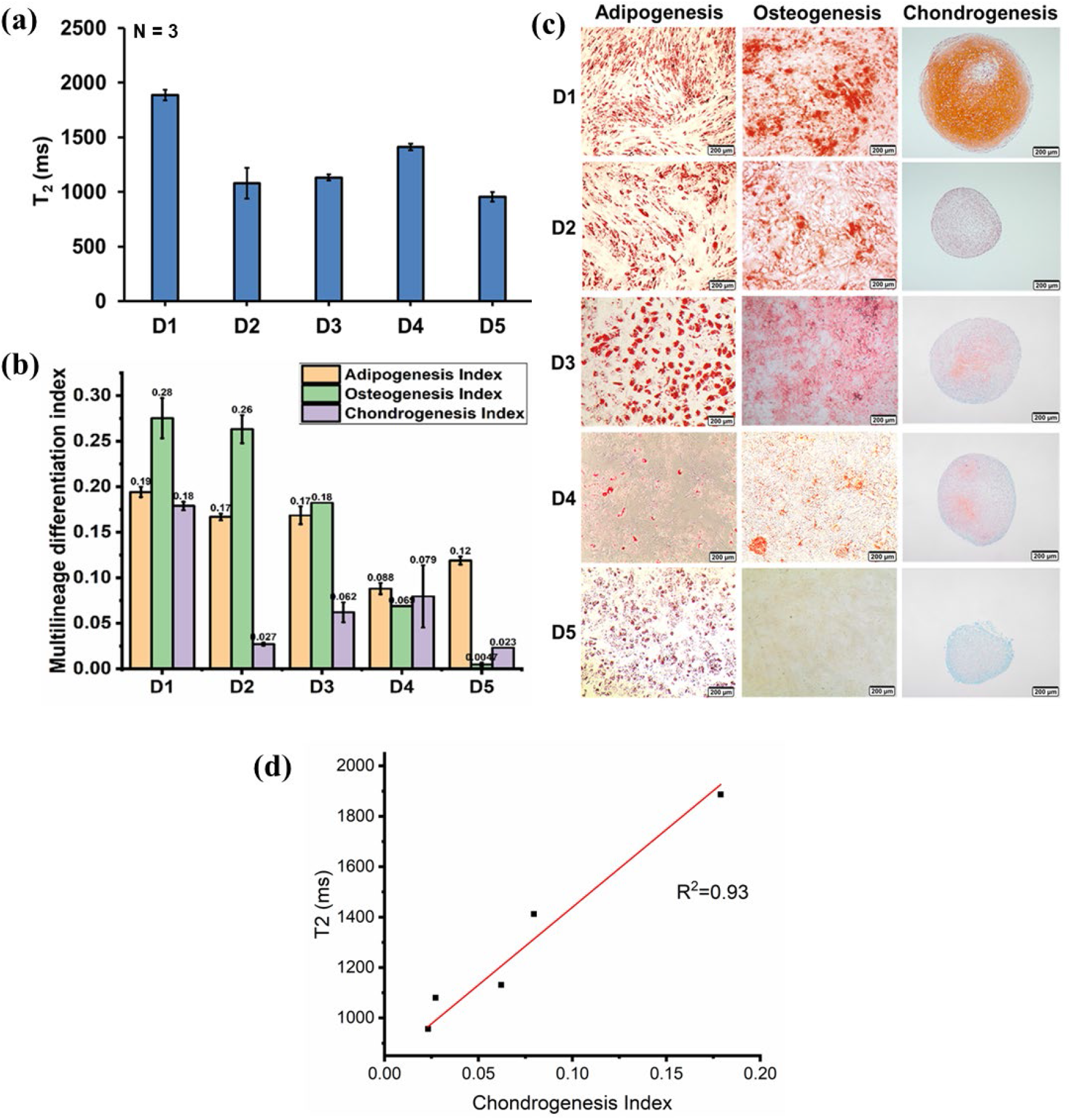
MRR analysis measures altered multi-lineage differentiation in MSCs from different passages and donors. (a) T_2_ values (n=3) of MSCs from different donors D1 to D5 (passage 3). (b) Image_based quantification of multi-lineage differentiation potential. Differentiation index (Adipogenesis Index, Osteogenesis Index, and Chondrogenesis Index) quantified from the images of respective MSC donors in passage 3. (c) Multilineage differentiation (Adipogenesis, Osteogenesis, and Chondrogenesis) images of MSCs from different donors.(d) T2 vs chondrogenesis index from different donors D1 to D5 showing T2 correlates well with condrogenesis potential (R2=0.93).

### MRR shows serial passage leads to an increase in the proportion of senescent MSCs

During *in vitro* culture of MSCs, cellular senescence tends to increase during serial passaging. Senescent cells can affect the proliferation of MSCs, differentiation potential, and hence the quality of cells for their clinical use. Therefore, it is important to identify senescent cells and potentially remove them from each passage of MSC culture.(Yin et al., 2018) We next analyzed MSCs (donor 4) from serial passages P3 to P8 under *in vitro* culture conditions. Using µMRR, T_2_ values of normalized concentrations of MSCs from each passage are shown in **Figure 6a**. We observed the highest T_2_ value for MSCs in the earliest passage, whereas these levels progressively decreased at later passages. MSCs from the later passage, where the Senescence Index was greater based on the intensity and number of cells displaying β-gal staining, showed the lowest T_2_ value (**Figure 6b and Figure S6**). Similarly, RT-qPCR of the cell cycle inhibitors p16 and p21 showed a trend toward highest expression in cells from late passages as compared to early passages (**Figure 6c**). Together, our data demonstrate the MRR T_2_ measurement as a non-destructive and label-free method for detecting the accumulation of senescent MSCs across a range of conditions and donors in live cultures.

**Figure 6.**
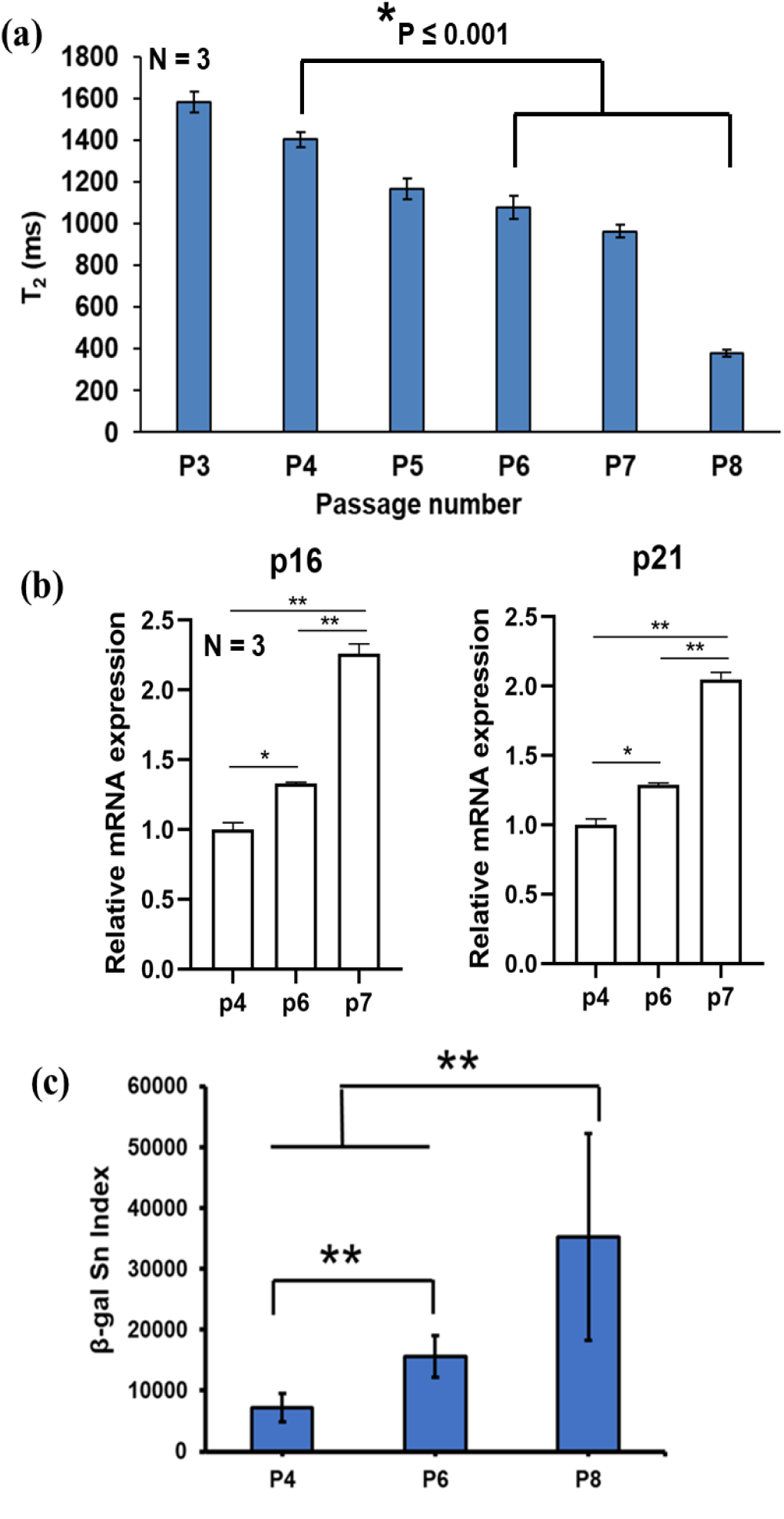
MRR analysis shows serial passage leads to an increase in the proportion of senescent MSCs. (a) Average T_2_ values (n=3) of MSCs which are passaged serially from 3 to 8. Statistical significance is determined by a two-tailed T-test. (b) Relative mRNA expression values of P21 and P16 for MSCs from the same donor for passage P6 and P7 compared to passage P4 as a control. The experiment was performed in technical triplicates. Unpaired two-tailed t-test was performed between selected pairs to determine the statistical significance. * represents P≤0.05, ** represents P≤0.01. (c) Quantification of senescence of β-galactosidase stained MSCs from the same donor for passage P4, P6, and P8. Statistical significance is determined by a two-tailed T-test, ** indicates p < 0.001. Ten to fifteen random areas were captured at the bright field by a color camera for quantifying the β-galactosidase staining-based senescence index (β-gal Sn Index) using MATLAB image processing.

### MRR measurements correlate with intracellular iron (Fe^3+^) and senescent level of MSCs

Because evidence suggests that senescent cells show an accumulation of iron (Fe^3+^), we next investigated the correlation between T_2_ values and Fe^3+^ levels. Fe^3+^ was quantified by a reversible fluorescent Fe^3+^ sensor (RPE)(Y. Wei; Aydin Z.; Zhang Y.; Liu Z.; Guo M., 2012) at the different passages of MSCs (D6, P4 vs. P7) in parallel with µMRR measurements. As shown in Figure **7a**, MSCs at the P7 show a statistically significant decrease in T_2_ value (P<0.001, 2-tailed T-test) compared to earlier passage P4, correlating with an increase of classical senescence markers (**Figure 7b and 7d**). **Figure 7c** shows that later MSC passages exhibit higher Fe^3+^ levels (*i*.*e*., higher Fe^3+^ mean intensity of 10k cells) as quantified by the RPE using flow cytometry and also with the brighter fluorescent intensity of cells in the images (**Figure 7e**). These results show the MRR measurement correlates with the Fe^3+^ accumulations in MSCs that also show a senescence phenotype.

**Figure 7.**
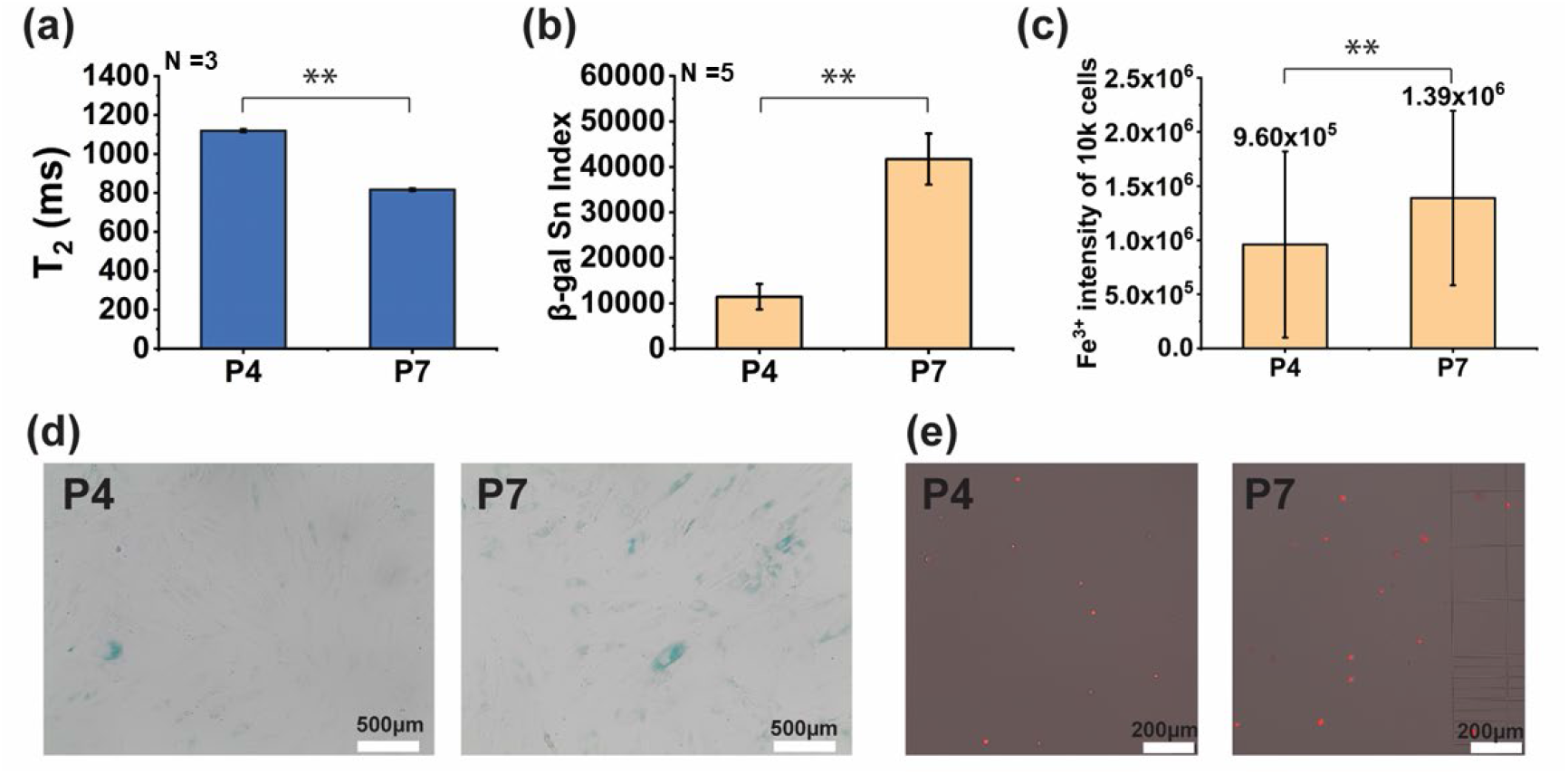
MRR measurements correlate with intracellular iron (Fe^3+^) and senescent level of MSCs. (a) MRR measurements, (b) senescence, and (c) Fe^3+^ quantification of donor 6 for passages P4 and P7. (d) Representative images of β-galactosidase staining of MSCs for passages P4 and P7 where the cytoplasm of senescent MSCs was stained as blue. Those images (images number n=5 for both 2 biological replicates; cell total number > 1K was used to quantify the β-gal Sn Index in Fig. b. (e) Representative images of Fe^3+^ staining of MSCs, the more senescent P7 cells showing the brighter intensity compared with the cells from earlier passage P4. Intracellular iron (Fe3+) level of 10K Fe^3+^ staining of MSCs was measured by flow cytometry (Fig. c). Statistical significance is determined by a two-tailed T-test, ** indicates p < 0.001.

## Discussion

We have established a rapid and non-destructive method for quantifying senescent cells *in vitro*. Our approach requires a small number of cells (<10^5^) and does not require any reagents or sample preparation steps. Direct Fe^3+^ quantification by µMRR provides well-correlated results with conventional end-point and time-intensive assays for senescence, such as RT-qPCR, Luminex assay, and β-galactosidase staining. Since paramagnetic Fe^3+^ (but not diamagnetic Fe^2+^) increases the magnetic susceptibility, which stimulates proton nuclear relaxation of water molecules in the cells, our measurement is specifically quantifying the Fe^3+^ content of cellular iron. Previously, intracellular Fe^2+^ (labile iron) was measured by colorimetric and other assays(Hirayama and Nagasawa, 2017), yet reliable and quantitative detection of Fe^2+^ has generally been challenging(Abbasi et al., 2021; Tenopoulou et al., 2007), presumably due to the reactive nature of Fe^2+^. For Fe^3+^ quantification, ferritin is often used as a surrogate maker(Mackenzie et al., 2008), which is a large (∼480kDa) protein complex that can accommodate up to 4500 atoms of Fe^3+^ iron. Thus, ferritin concentration may not correlate well with the Fe^3+^ amount stored in the cell. ICP-MS quantification(Denoyer et al., 2016) of total iron concentration measurement is the current standard for iron quantification; however, this metric may not correctly reflect the stored Fe^3+^, which is more closely correlated to cellular senescence.

In contrast to the conventional iron quantification methods, µMRR is uniquely positioned to advance iron biology in general. µMRR allows phenotyping of live cells, allowing downstream biological and functional measurements of other indicators on the same cells. This ability enables direct correlations between MRR T_2_ and other biochemical and functional measurements. Thus, adapting µMRR is potentially transformative for the field of cell therapy production, especially given the lack of consensus cell surface markers specific for senescent cells.

In this study, the multilineage differentiation potential of MSCs from different donors also correlates well with MRR results, with the highest T_2_ value measured from MSCs having robust multilineage differentiation potential. This technology will be instrumental for screening MSCs for the highest growth and differentiation potential in a patient-specific manner in real-time. One can also adopt this new bioengineering modality for optimizing the biomanufacturing of MSC or other cell therapy products. For example, MRR could be used to monitor cell cultures in real-time in a way that would allow the removal of senescent cells using microfluidic sorting(Yin et al., 2018). Thus, our approach could overcome a major bottleneck to produce higher grade MSCs and cell products for specific therapeutic modalities. In addition, recent data suggest important roles of iron homeostasis in the maintenance of iPSCs(Han et al., 2019),(Han et al., 2020) and HSCs(Xin Jin et al., 2018),(Muto et al., 2017), raising the prospect that this technology can impact a broader class of therapeutic cells in the future.

More broadly, the ‘iron-imaging’ modality utilized here is already widely used in T_1_ and T_2_ weighted imaging using MRI (Dusek et al., 2013; Golay et al., 2001; Parikh et al., 2008; Van Walderveen et al., 2001). Therefore, any insights from MRR quantification of cellular senescence could readily be compared to *in vivo* pathology. Given the resolution of modern MRI approaches below 100µm(Stucht et al., 2015), non-invasive detection of senescent cells in live tissue may be feasible in the future. We envision µMRR as a rapid, live-cell quality screening tool for detection fo cellular senescence across a range of cell types and tissues.

### Experimental procedures

#### MSC Culture

Bone marrow-derived mesenchymal stem cells (MSC) were purchased from Lonza Pte. Ltd. and Rooster Bio Inc. The cells were expanded in tissue culture plate (TCP) at an initial cell density of 1500 cells/cm^2^ in low glucose Dulbecco’s Modified Eagle Medium (DMEM) supplemented with 10% fetal bovine serum (FBS), 1% GlutaMAX, and 1% Penicillin/ Streptomycin (Thermo Fisher Scientific, Singapore) at 37°C in 5% CO_2_ atmosphere. The medium was changed every two days, and the cells were harvested at 80% confluency for further experiments or subcultures. During the cells harvesting, the cells were incubated with 0.25% Trypsin-EDTA (Thermo Fisher Scientific, Singapore) for 3 minutes. The cell number was calculated using a disposable hemocytometer (INCYTO, Korea) with the trypan blue exclusion method. MSCs at passages 3 (P3) to 6 (P6) were used for further experiment unless otherwise stated. MSC senescence induction was carried out by treatment with culture media containing 10 or 20 ng/ml of TGF-β1 for three days. For doxorubicin (DOX) treatment, cells were seeded at 1500 cells/cm^2^ and incubated with 1µM doxorubicin (Sigma) for 24 hours at 37 °C. Subsequently, the medium was removed, and a fresh medium was added to the cells to incubate for 24 hours before analysis.

#### MSC sorting with inertial spiral microchannel device

The inertial spiral microchannel device was designed and fabricated in the same way as previously described.(Yin et al., 2018) The device has 8 loops with a radius decreasing from 12mm to 4mm and a trapezoidal cross-section with 580µm width, 85um inner, and 133 µm outer height. It has one inlet for the introduction of the cell suspension to be sorted and two outlets for the collection of sorted cells. The design was carved on a micro-milled aluminum mold (Whits Technologies Inc., Singapore), and the device was cast from polydimethylsiloxane (PDMS) with a ratio of 10:1 base and curing agent mixture (Sylgard 184, Dow Corning Inc., USA).

Before sorting, MSC was resuspended in culture media at 1-2 million cells/ml and loaded into syringes (Thermo Fisher Scientific, Singapore) that were connected to Tygon tubing (Spectra Teknik Pte. Ltd., Singapore). The tubing was inserted into the inlet of the device, and two separates tubing was inserted at the outlets for the collection of sorted cells. A syringe pump (PHD2000, Harvard Apparatus Inc., USA) was used to control the flow rate of cell suspension in the device. Two serials sorting was performed to separate MSC into three populations of different sizes. The first sorting was carried out at 3.5ml/min, and the cells collected at the inner outlet are the largest subpopulation (22-26 µm). The cells collected at the inner outlet were subjected to second sorting at 1.5ml/min, and the cells collected at the inner and outer outlets are medium size subpopulation (15-22µm) and smallest subpopulation (11-15µm), respectively. The unsorted MSC was used as a control in this experiment. Real-time visualization of the separation within the inertial spiral microchannel device was achieved by an inverted microscope (IX71, Olympus Co., Japan) equipped with a high-speed CCD camera (Phantom v9, Vision Research Inc., USA).

#### MRR measurement

MRR consists of a portable, permanent magnet (Metrolab Instruments, Plan-les-Ouates, Switzerland) with *B*_0_ = 0.5 T and a bench-top type NMR console (Kea Magritek, Wellington, New Zealand). ^1^H MRR measurements were performed at the resonance frequency of 21.65 MHz inside the magnet. A single resonance proton MRR probe with a detection micro-coil of 900-μm inner diameter was used for accommodating the MRR samples into the microcapillary tubes (o.d.: 1,500 μm, i.d.: 950 μm) (22-260-950, Fisherbrand, Waltham, MA, USA). In the MRR probe, the electronic parts and coil were mounted on the single printed circuit board (**Figure 1c**).(Peng et al., 2014) All the experiments were performed at 26.3°C inside the magnet maintained by a temperature controller (RS component, UK).

For all MRR experiments, the normalized concentration (60K cells in 4µL volume) of MSCs have been used unless otherwise stated. The MSCs samples were spun down at 300 g for 5 minutes, and the supernatant was aspirated. The pellet is suspended in 20µl of PBS and filled at a 4mm length of the micro-capillary tube. The micro-capillary tube was sealed with critoseal (Leica Microsystems) and mounted into the coil for MRR measurements (**Figure S2**). Proton transverse relaxation times (T_2_) were measured by standard Carr-Purcell-Meiboom-Gill (CPMG) pulse programme(Carr and Purcell, 1954; Meiboom and Gill, 1958) (**Figure 1b**). We maintained the transmitter power output at 12.5 mW for a single 90° pulse of pulse length 6μs for all the T_2_ measurements. The CPMG train of pulses with inter echo time of 200μs with 4000 echoes was used for all experiments. A recycle delay of 2 s, which is sufficient to allow all the spins to return to thermal equilibrium, was used. 24 scans were performed for all experiments for signal averaging.

#### Real-time polymerase chain reaction (RT-qPCR) analysis

Total RNA was extracted with the RNeasy Mini Kit (Qiagen, USA) following the manufacturer’s protocol. The concentration of RNA was determined using a NanoDrop UV-vis Spectrophotometer (NanoDrop Technologies, USA). Then, reverse transcription reaction was carried out with 200ng total RNA using iScript™ cDNA synthesis kit (Biorad Laboratories, USA). Real-time PCR was performed using SYBR Green system with primers listed in **Table 1**. ABI 7500 real-time PCR system (Applied Biosystem, USA) was used to performed real-time PCR at 95°C for 10mins and 40 cycles of amplification which encompasses denaturation step at 95°C for 15s and extension step at 60°C for 1min. The gene expression level was normalized to glyceraldehyde-3-phosphate dehydrogenase (GAPDH) and calculated using the 2^-ΔΔCt^ formula with reference to the respective control group. The data presented is representative of 3 technical replicates for each condition.

**Table 1.**
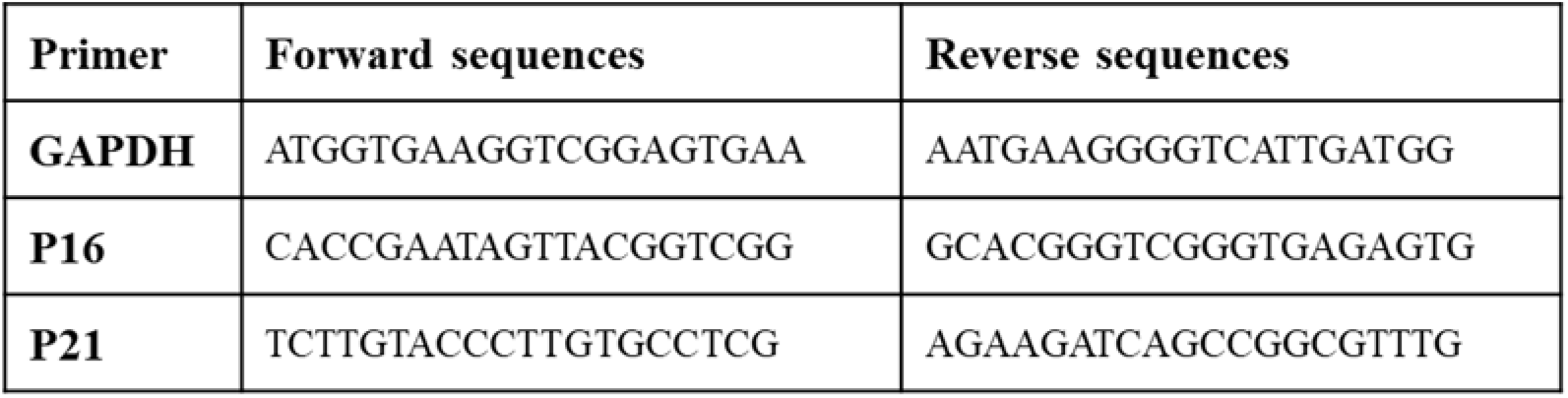
The primers for Real-time PCR

**Table 2.** Description of Donors

| **Name** | **Source** | **Age** | **Sex** |
| --- | --- | --- | --- |
| Donor 1 | Human MSC | 20 Y | M |
| Donor 2 | Human MSC | 22 Y | M |
| Donor 3 | Human MSC | 25 Y | M |
| Donor 4 | Human MSC | 19 Y | F |
| Donor 5 | Human MSC | 24 Y | M |
| Donor 6 | Human MSC | 36 Y | M |

#### Conditioned media and ELISA

Size-sorted MSCs were seeded in T75 flasks for overnight adherence before media removal the next day. The cells were washed with PBS, and cultured in serum-free media for 48 hours at 37°C, with 5% CO_2_ concentration. The media was then collected and centrifuged at 4500g for 15 minutes to remove cell debris. The supernatant was collected and concentrated to 500µl with an Amicon Ultra 15 filter (3 kDa cut-off membrane) and stored at −80°C before analysis. Cell number in each condition was determined to normalize the concentration values of the analytes of interest (MCP-1, IL-6, IL-8, and TGF-β1) obtained using Luminex ®-based multiplex assays (R&D Systems) according to manufacturer’s instructions. All samples were analyzed in technical duplicates and performed for two different donors. The data were obtained with a MAGPIX reader (Millipore), and concentrations were derived from measured mean fluorescence intensities (MFI) using fitted standard curves using 5-parameter logistic regression (SSL5) using the Milliplex Analyst software (Millipore).

#### *In vitro* multilineage differentiation assay and quantification

MSCs were seeded in 24-well plates for overnight adherence before media removal and replaced with specific differentiation media (STEMCELL Technologies) for osteogenesis and adipogenesis, following the manufacturer’s protocol. Osteogenesis induction was confirmed with Alizarin Red S staining (ScienCell) for calcium deposits, while adipogenic differentiation was determined with Oil Red O staining (Sigma) for the detection of lipid droplets. For chondrogenesis, MSCs were pelleted at 1×10^6^ cells/pellet in a 15ml tube at 300 g, 5 minutes. Chondrogenic media (STEMCELL Technologies) was added to this pellet, and media was changed every alternate day for three weeks before glycosaminoglycan staining with Safranin O.

The multilineage differentiation index (*i*.*e*., Adipogenesis Index, Osteogenesis Index, and Chondrogenesis Index) was computed as the product of the relative stained area and the relative red color intensity of the stained area in an image. The positive staining areas were extracted by the adaptive threshold algorithm(Yang et al., 2017) developed using MATLAB software (MathWorks, USA). The relative stained area was calculated with reference to the image area. The relative red color intensity was calculated with reference to the maximum intensity in the RGB image.

#### Senescence assay and quantification

MSCs were seeded on a 6-well plate at an initial cell seeding density of 1500 cells/cm^2^ and cultured for 3-5 days. Senescence β-galactosidase staining kit (Sigma Aldrich) was used to identify the senescent cells according to the manufacturer’s protocol. Briefly, the cells were fixed at room temperature for 10mins and incubated in a staining solution containing β-gal at 37°C overnight. Cells stained positive with blue precipitate as a result of β-gal substrate cleavage were indicative of senescent cells.

Five to fifteen random areas (except for the Figure 3b) were captured at the bright field by a color camera for quantifying the β-galactosidase staining-based senescence index (β-gal Sn Index) using MATLAB image processing. The areas of cells stained positive were green/blue color and reflected as the areas lacking red color. Thus, the positive staining areas were extracted by the adaptive threshold algorithm(Yang et al., 2017) processing at the red channel of RGB images. β-gal Sn Index was computed as the total sum of the complement of red intensity (mean of the red intensity of the image – red intensity) in all selected areas. The final β-gal Sn Index was normalized by the confluency of cells.

#### Fe^3+^ staining and quantification of MSCs

A reversible fluorescent Fe^3+^ sensor (RPE)(Y. Wei; Aydin Z.; Zhang Y.; Liu Z.; Guo M., 2012) was used to stain the intracellular iron (Fe^3+^) of MSCs. A stock solution of RPE (1 mM in acetonitrile) was diluted to a concentration of 20 µM in PBS for Fe^3+^ staining. Cells were washed twice in PBS and incubated with PBS containing RPE at 37°C for 30 minutes. After incubation, the cells were washed twice and suspended in PBS for flow cytometry measurement and microscope imaging.

#### Immunofluorescence and confocal microscopy

For detection of SA-βgal together with another senescence marker (γ-H2AX foci)(Bernadotte et al., 2016) and the proliferation marker, 5-ethynyl-2′-deoxyuridine (EdU) in the cells, we performed the protocol as previously mentioned(Itahana et al., 2013). Briefly, the cells were incubated with 10µM EdU (Click-iT® EdU Alexa Fluor® 488 Imaging Kit, Invitrogen) for 24 hours before fixation with 10% neutral buffered formalin (Sigma) and β-gal staining (Sigma Aldrich) using components from the staining kit according to the manufacturer’s instructions (#9860, Cell Signaling Technology). Subsequently, EdU labeling was carried out following the manufacturer’s protocol (Invitrogen).

For immunostaining, cells were washed in PBS and incubated with blocking buffer (PBS containing 0.5% bovine serum albumin) for 1 hour before incubation with a mouse anti-phospho-Histone H2A.X (Ser139) antibody (05-636, Millipore) in blocking buffer for overnight at 4°C. Cells were then incubated with Rhodamine Red™-conjugated secondary antibody (Jackson Immuno Research Laboratories) for 1 hour, washed thrice with PBS, and counterstained with NucBlue (Hoechst 33342)-containing mounting solution (Invitrogen). Images were acquired with an FV1200 Confocal Microscope using a 20x objective lens (Olympus). β-gal-stained cells were also visualized under phase-contrast microscopy.

#### Statistical analysis

Statistical analyses were performed using Microsoft Excel and Origin pro (Figure 7). All data were presented as the mean ± SD Groups were compared using the two-tailed t test. A p value <0.05 was considered statistically significant.

## Acknowledgments

This research is supported by the National Research Foundation, Prime Minister’s Office, Singapore under its Campus for Research Excellence and Technological Enterprise (CREATE) programme, through Singapore-MIT Alliance for Research and Technology (SMART): Critical Analytics for Manufacturing Personalized-Medicine (CAMP) Inter-Disciplinary Research Group.

## Author Contributions

S.S.T, L.A.B., and J.H. designed the overall experimental plan. S.S.T. carried out the MRR experiments with support from C.A.T, S.H.N, and D.Y., C.A.T., S.H.N., and R.O. performed various biochemical assays. S.S.T. and D.Y. performed data analysis. The manuscript was written by S.S.T. and all authors participated in editing and revising the manuscript.

