## Supplementary Information for "Rapid and Live-cell Detection of Senescence in Mesenchymal Stem Cells by Micro Magnetic Resonance Relaxometry"

MSC is seeded on
cell culture flask in expansion medium

MSCs were harvested at 80% confluency. Count the cell number

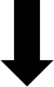

Normalize the cell number to 3×10^5^ cells in 1ml

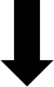

Spin down 200g 5 min, aspirate supernatant. Suspend the pellet into 20µl of PBS.

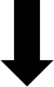

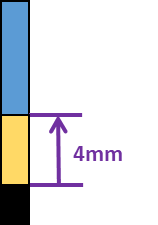

Fill 4 µl (6×10^4^ cells) into the microcapillary tube (4mm range, yellow color).

Seal the capillary with crystoseal (black color)

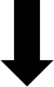

Place the sample capillary tube into the radio frequency detection coil of MRR magnet.

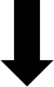

T_2_ measurements

Figure S1: MSC sample preparation workflow for T_2_ measurements.

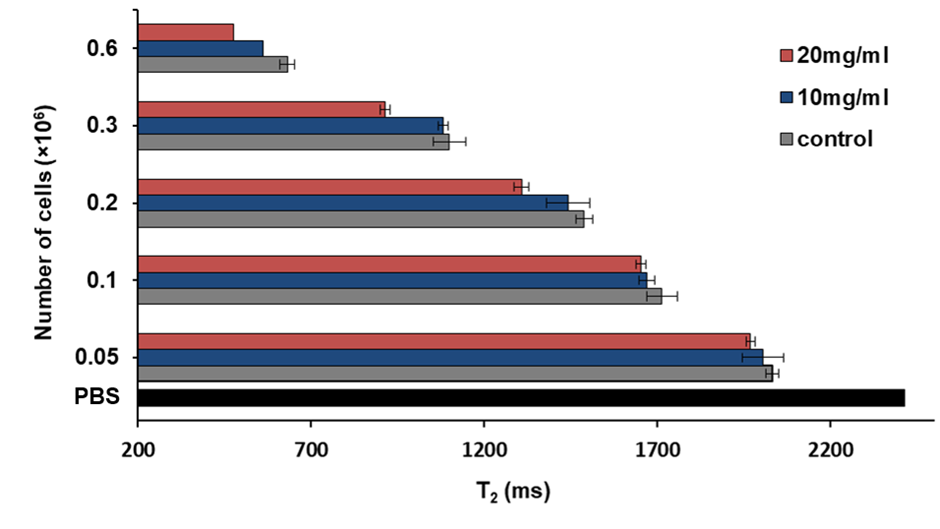

Figure S2: Limit of detection of MRR for senescent MSCs. T_2_ values of different cell concentrations of control and TGF-β1 (10 and 20 mg/ml) treated MSCs (Donor 3, passage 4). The T_2_ value of PBS is shown as a black bar at the bottom of the figure.

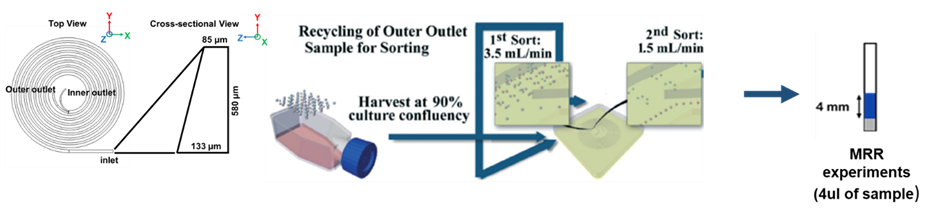

Figure S3: The protocol to separate senescent MSCs from culture expansion and MRR detection. The spiral microfluidic device, as shown in the figure, has one inlet and two outlets(Yin et al., 2018) . The MSC culture is pumped into the microfluidic device and sorted at a speed of 3.5ml/min to collect the Large cells (22-26µm) from the inner outlet and the cells from the outer outlet sorted again at a speed of 1.5ml/min to collect small and proliferating cells (11-15µm) and medium-sized cells (15-22µm). Normalized concentrations of all sorted cells are filled in a microcapillary tube and analyzed by MRR.

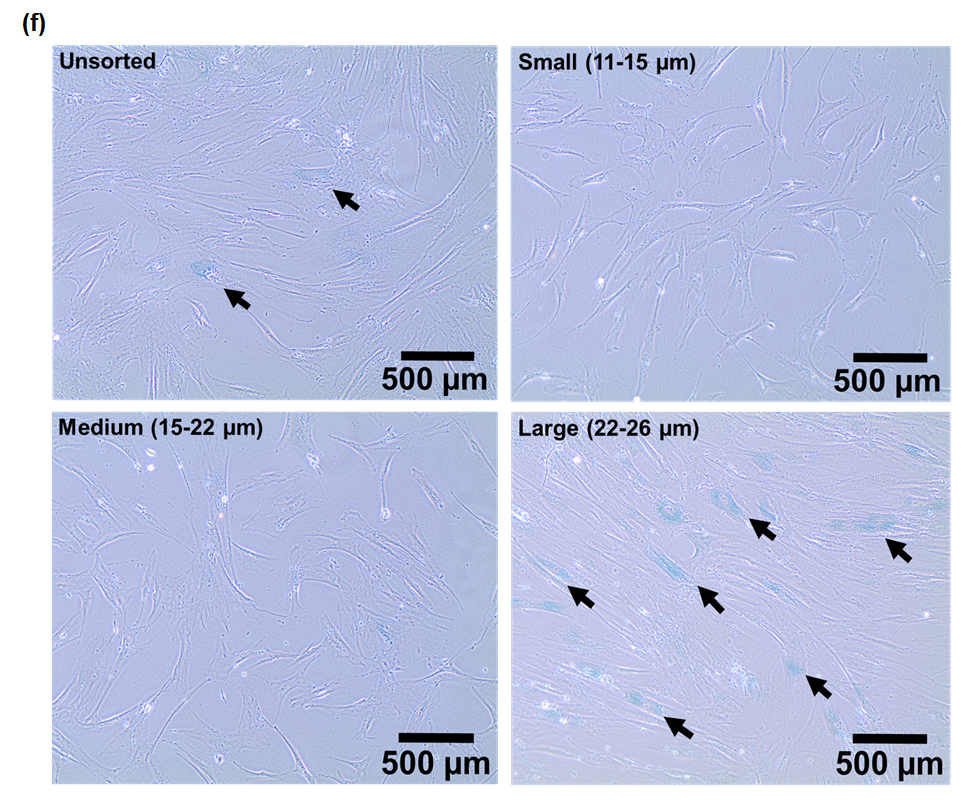

Figure S4: β-galactosidase staining of unsorted and size sorted MSCs where the cytoplasm of senescent MSCs are stained as blue (black arrows).

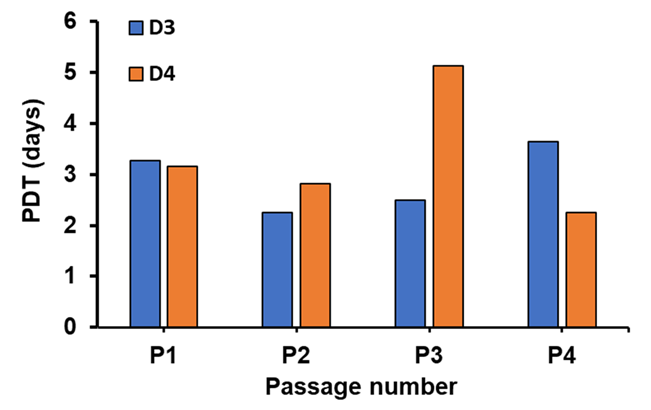

Figure S5: Population doubling time (PDT) of MSCs from different donors. PDT (days) of donors 3 and 4 from passage number 1 to 4.

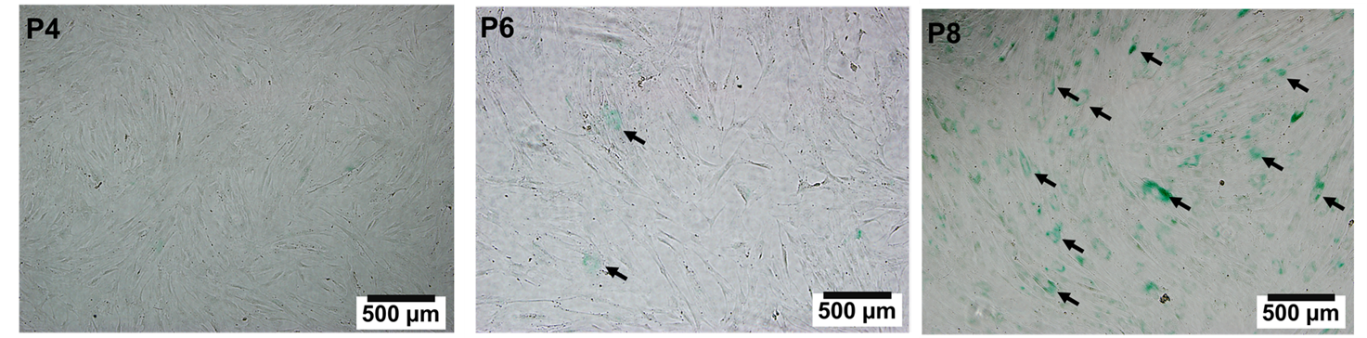

Figure S6: β-galactosidase staining of MSCs from the same donor for passage P4, P6, and P8 where the cytoplasm of senescent MSCs was stained as blue (black arrows).
